## Supplementary material for "Thermodynamic genome-scale metabolic modeling of metallodrug resistance in colorectal cancer"

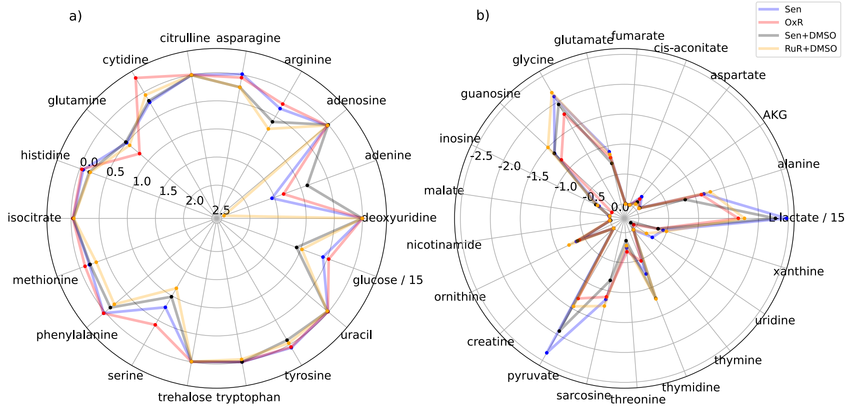


**Figure S1 Experimentally determined exchanged fluxes used to constrain the HCT116 - specific genome-scale metabolic model.** Metabolite concentrations in the standard and DMSO media were determined at four different time points (0, 24, 48, $72\,\mathrm{h}$) and were used to calculate an uptake rate ($\mathrm{mmol\,g^{-1}\,h^{-1}}$) as outlined in the Materials and Methods. Uptake rates (a) and excretion rates (b) are shown for HCT116 cells resistant to oxaliplatin (OxR, red) or ruthenium (RuR + DMSO, orange) and their sensitive controls a standard (Sen, blue) and a 0.5% DMSO medium (Sen + DMSO, black). The exchange fluxes for lactate and glucose were scaled down by 15, as indicated by the labels, to fit into the plot.

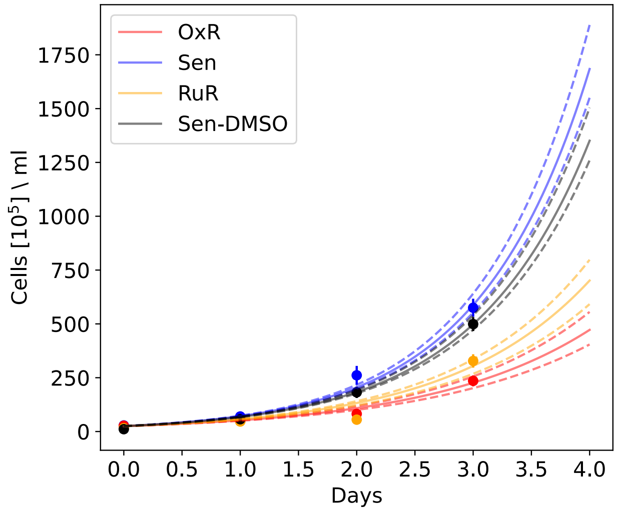


**Figure S2 Observed differences in growth rate between drug treatments and their sensitive (Sen) controls.** HCT116 cells resistant to oxaliplatin (OxR, red) or Bold-100/KP1339 (RuR + DMSO, orange) and their sensitive controls were inoculated in a standard (Sen, blue) and a 0.5% DMSO medium (sensitive + DMSO, black), respectively (see Materials and Methods for details). Cell density measurements were taken every 24 hours. The mean standard deviation of the three replicates measured at each time point is shown. Fitting an exponential growth curve to the mean values using the scipy package (Version 1.5.2) in Python (Version 3.7.9), the growth rate ($\mu$) was fitted to minimize the averaged sum of squared residuals for each condition (solid line). Growth curves fitted to the upper and lower standard error (dashed lines) are also shown and were used to set upper and lower bounds in the flux analysis.

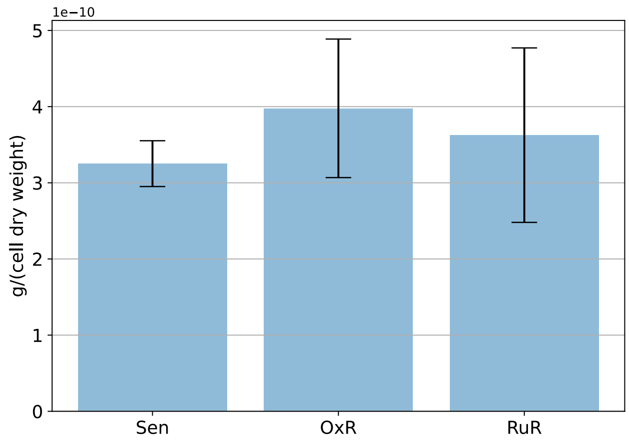


**Figure S3 Cell dry weight of wild-type and resistant cells.** Dry mass of HCT116 sensitive (Sen), oxaliplatin-resistant (OxR) and Bold-100/KP1339-resistant (RuR) cells were measured as outlined in the Materials and Methods. Mean standard errors of 4 biological replicates are shown.

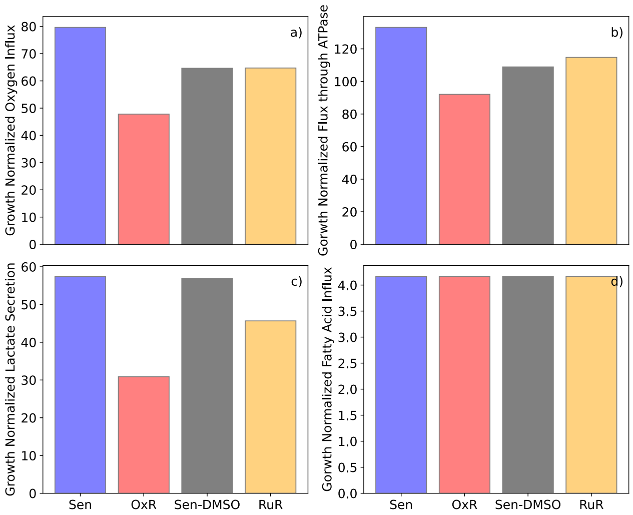


**Figure S4 Comparison of fluxes through key energy metabolism reactions for ruthenium- and oxaliplatin-based resistances before growth rate normalization.** Non-normalized flux values through (a) the oxygen uptake reaction - HMR_9048, (b) the ATP synthase reaction - HMR_6916, (c) the lactate secretion reaction - HMR_9135, and (d) the fatty acid influx - sum of m01362s_FAx, m02387s_FAx, m02389s_FAx, m02646s_FAx, m02674s_FAx, m02938s_FAx, across the four model instances (sensitive - Sen - blue bars; sensitive in a DMSO-based medium - Sen-DMSO - gray bars; oxaliplatin-resistant - OxR - red bards; Bold-100/KP1339 - RuR - yellow bars) are shown. Fatty acid influx is the combined influx of stearate, palmitate, oleate, linolenate, linoleate, arachidonate.

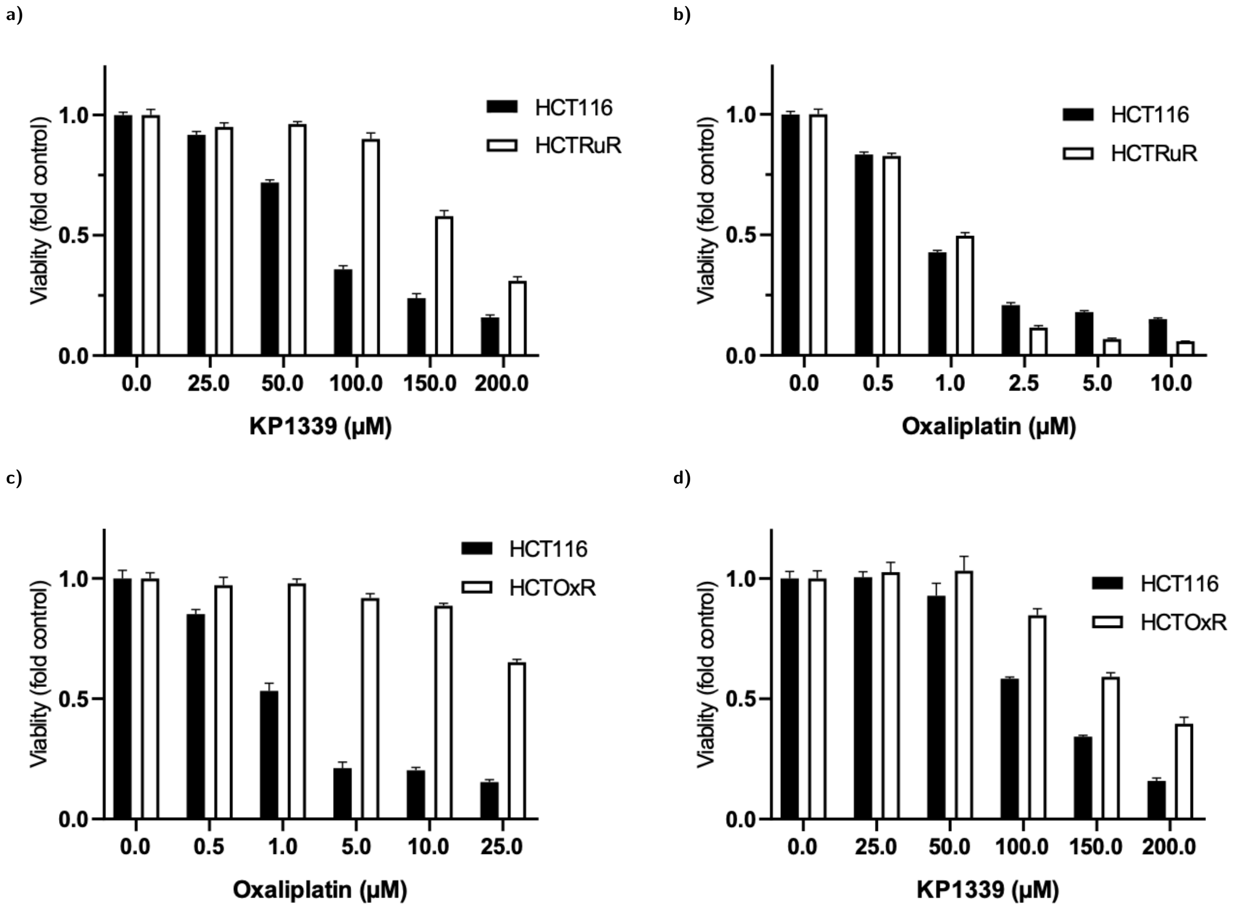


**Figure S5 Investigation of cross-resistance of the resistance model HCT116 with acquired BOLD-100/KP1339-resistance (RuR) towards oxaliplatin and cross-resistance of HCT116 with acquired oxaliplatin-resistance (OxR) towards BOLD-100/KP1339.** Viabilities are expressed in fold-change to untreated control in the function of the applied drug concentrations. (a) Parental sensitive HCT116 cells (HCT116) vs. RuR cells treated with the Ru-based BOLD-100/KP1339. (b) HCT116 vs. RuR cells treated with oxaliplatin. (c) HCT116 vs. OxR cells treated with oxaliplatin. (d) HCT116 vs. OxR cells treated with BOLD-100/KP1339.

### Mito stress test

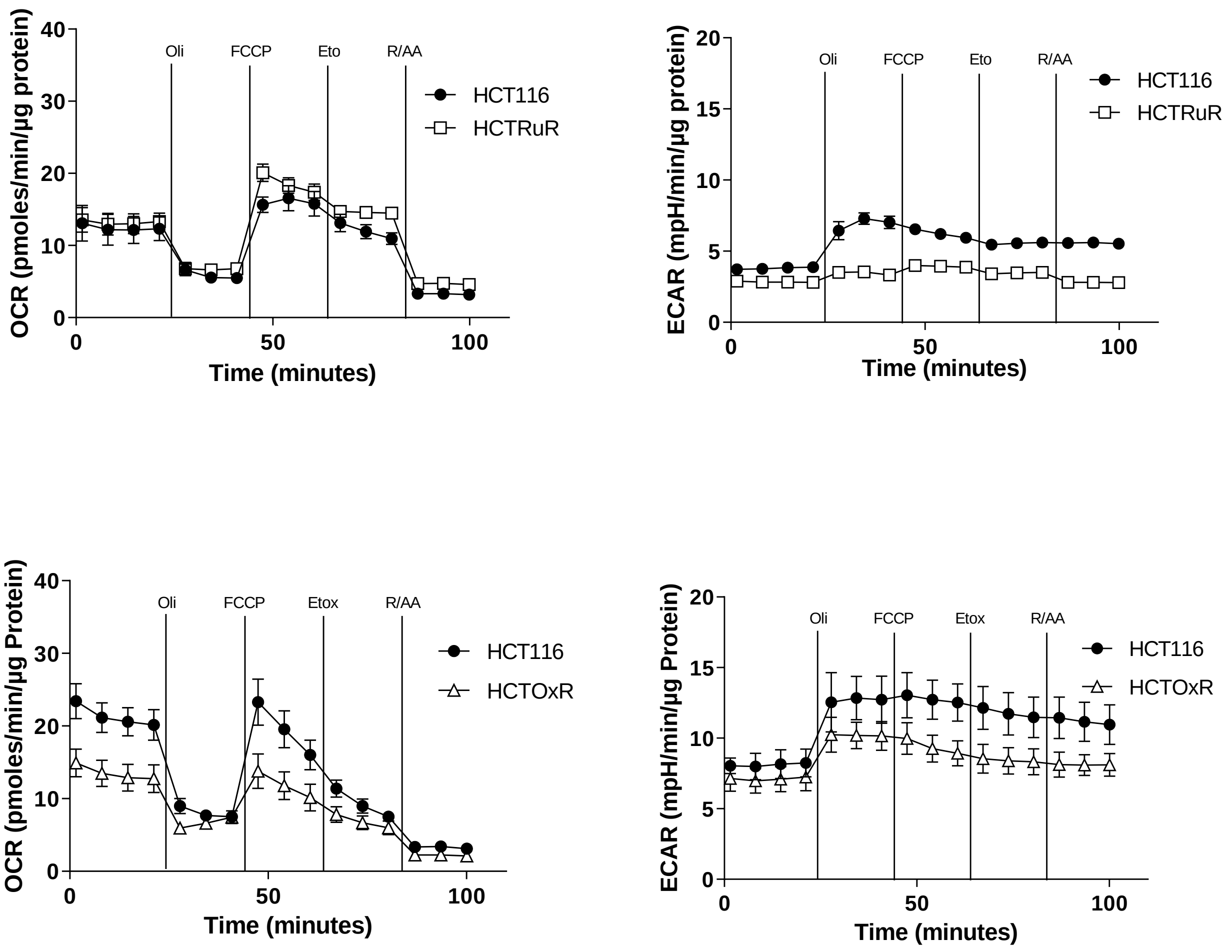


**Figure S6 Mito Stress test measuring the metabolic parameters OCR and ECAR with the Seahorse FX analyzer of sensitive vs. oxaliplatin-resistant HCT116 cells (HCT116 vs. HCTOxR); and sensitive vs. BOLD-100/KP1339-resistant HCT116 cells (HCT116 vs. HCTRuR).** The points and error-bars correspond to mean ± SD for each group ($n = 3$). Oxygen consumption rate (OCR) on the left and extracellular acidification rate (ECAR) on the right of the panel. All values have been normalized to total protein content in $\mu$g.
